## Supplementary figures for "CodonTransformer: a multispecies codon optimizer using context-aware neural networks"

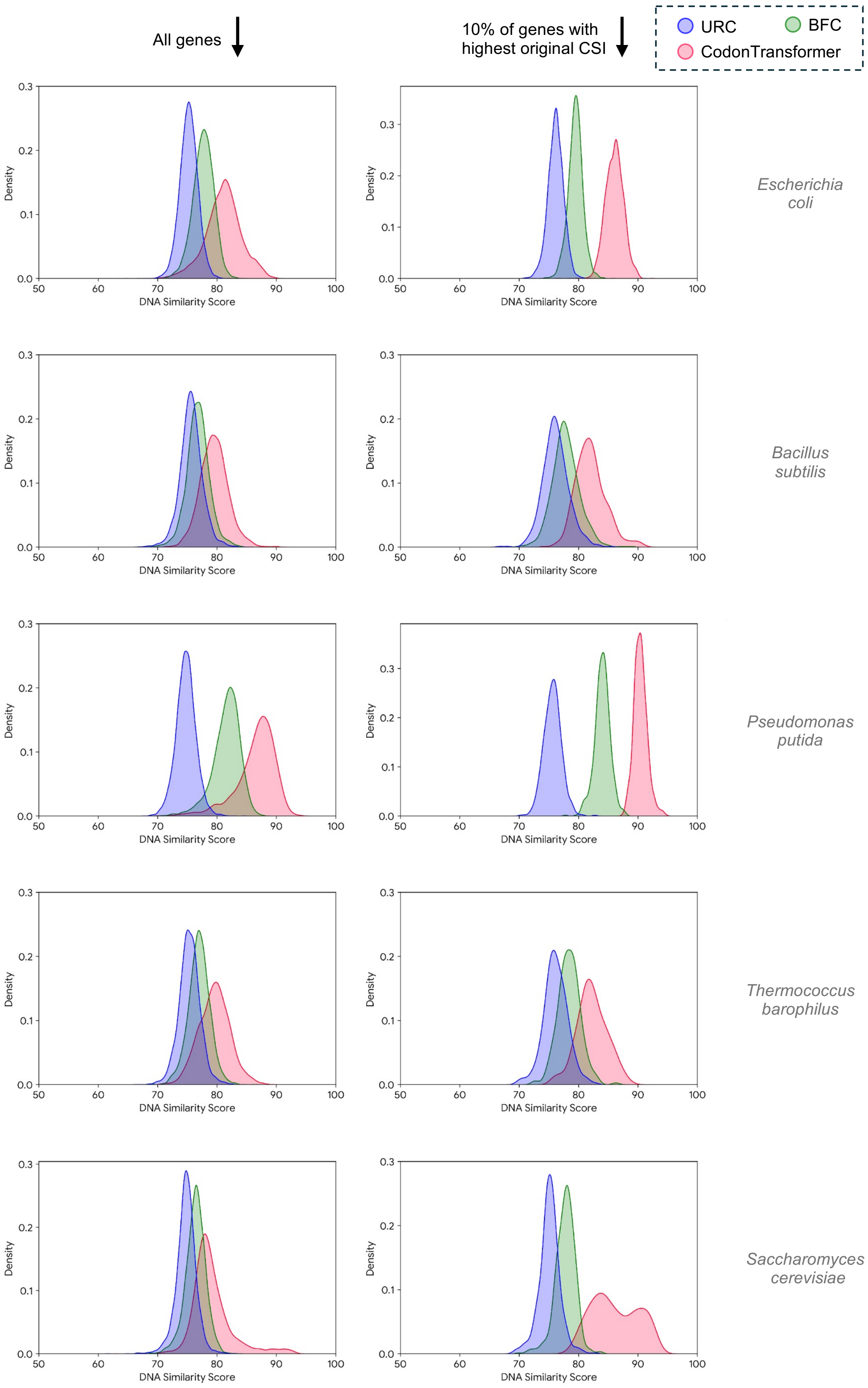


**Supplementary Fig. 1, continued on the next page.**

**Continued Supplementary Fig. 1**

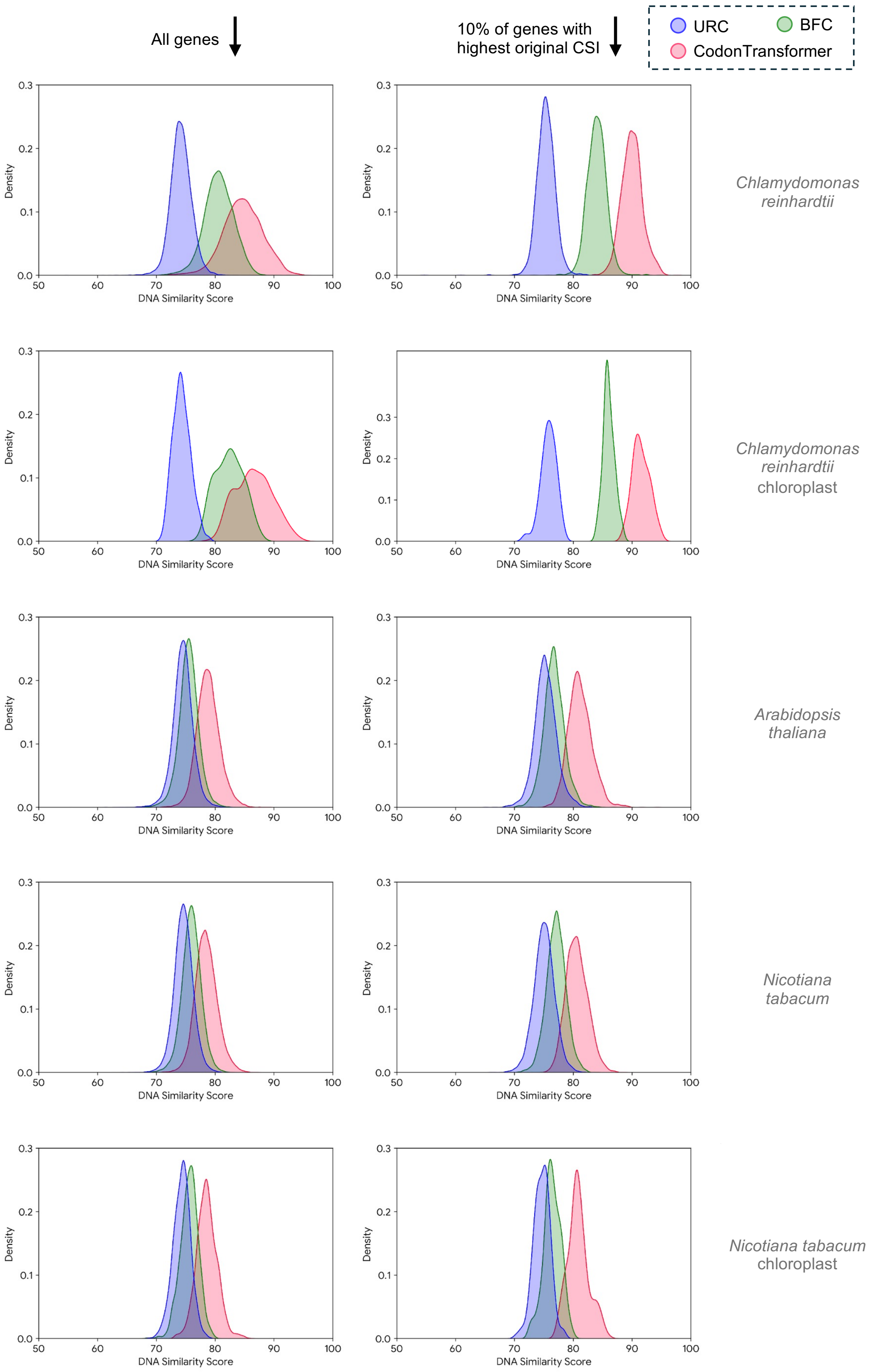


**Supplementary Fig. 1, continued on the next page.**

**Continued Supplementary Fig. 1**

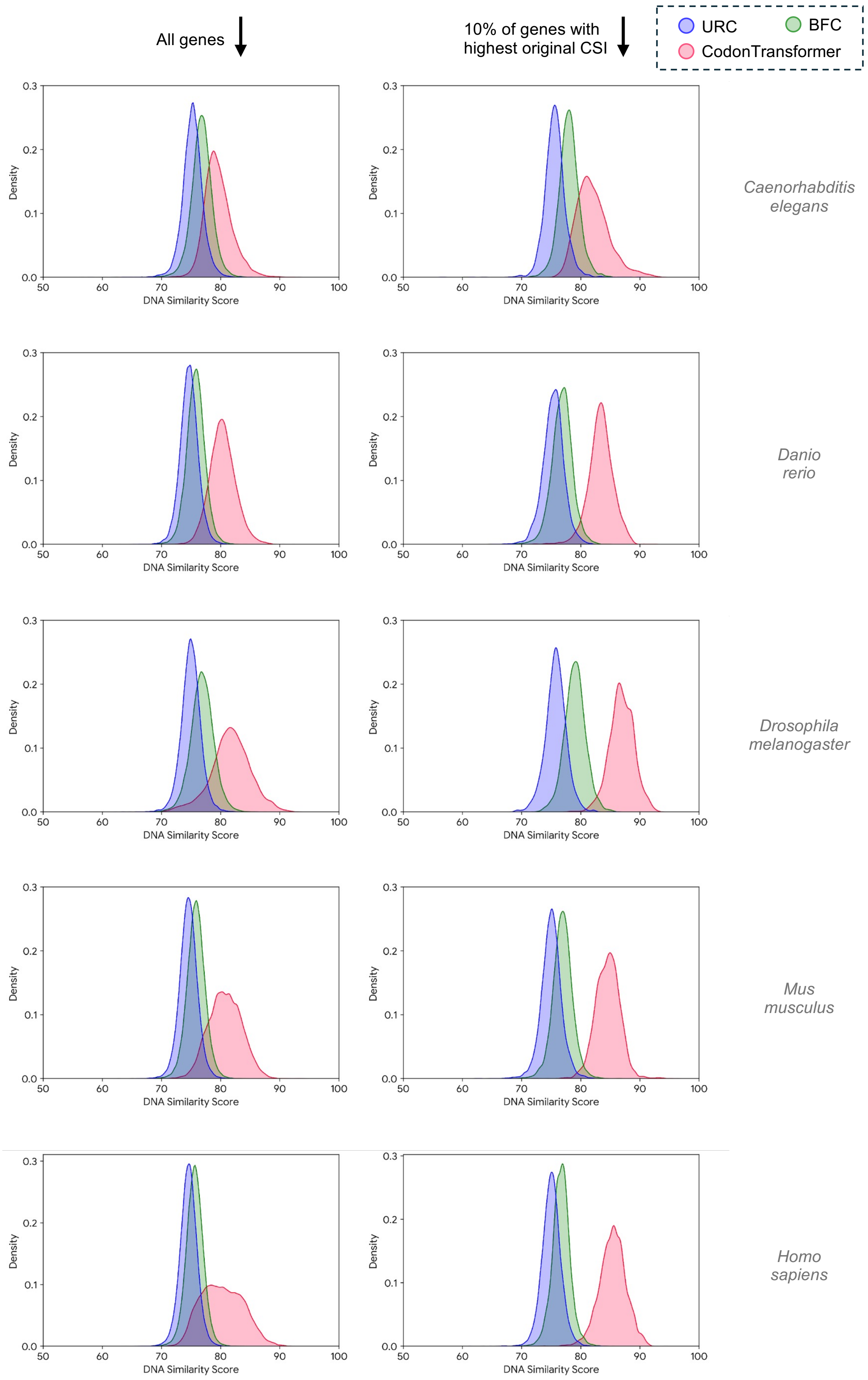


**Supplementary Fig. 1:** Kernel density plots for DNA similarity between original genes and DNA sequences designed by CodonTransformer (red), codons with Unified Random Choice (URC, blue) and Background Frequency Choice (BFC, green). The left plots are for all genes for each organism and right plots are for 10% genes with the highest original CSI).

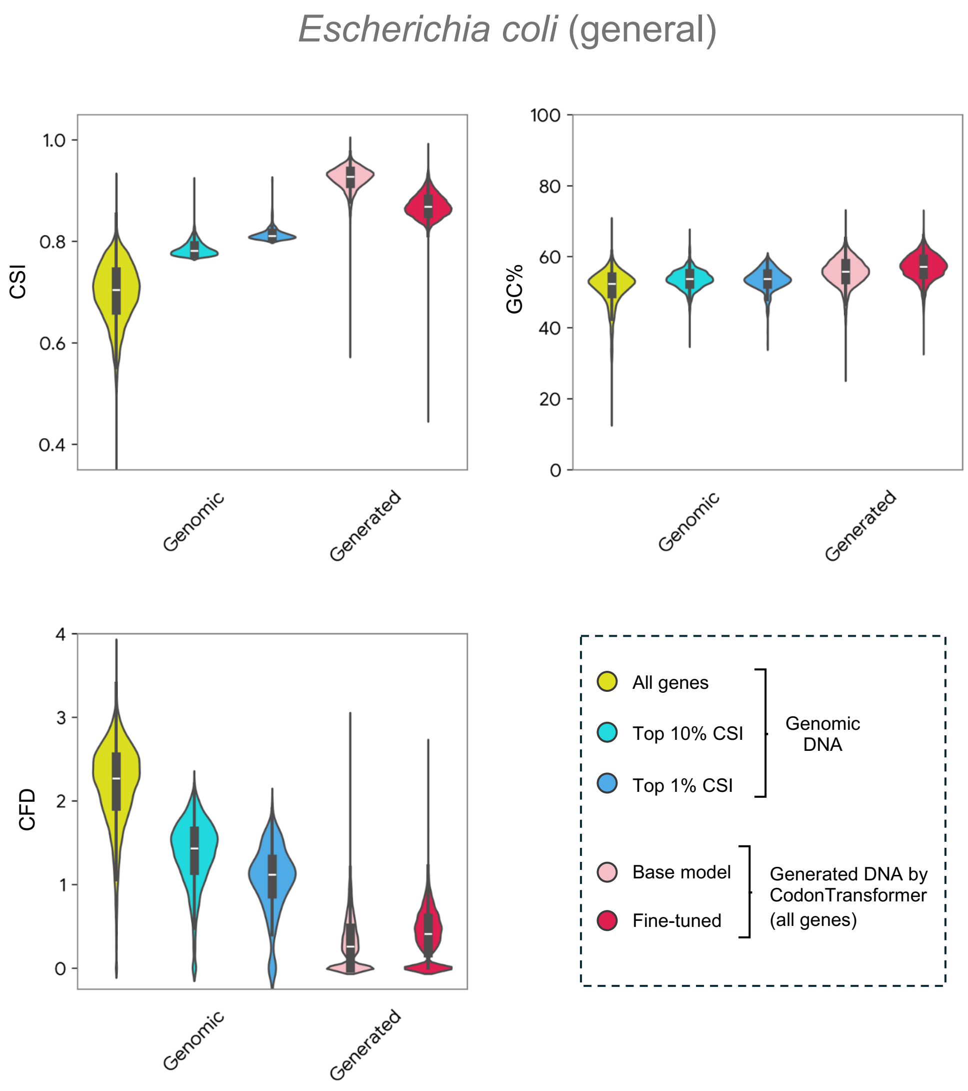


**Supplementary Fig. 2:** Codon similarity index (CSI), GC content, and codon frequency distribution (CFD) of genomic DNA sequences of *E. coli* general (merged *E. coli* genomes) and their generated counterparts by the base and fine-tuned CodonTransformer.

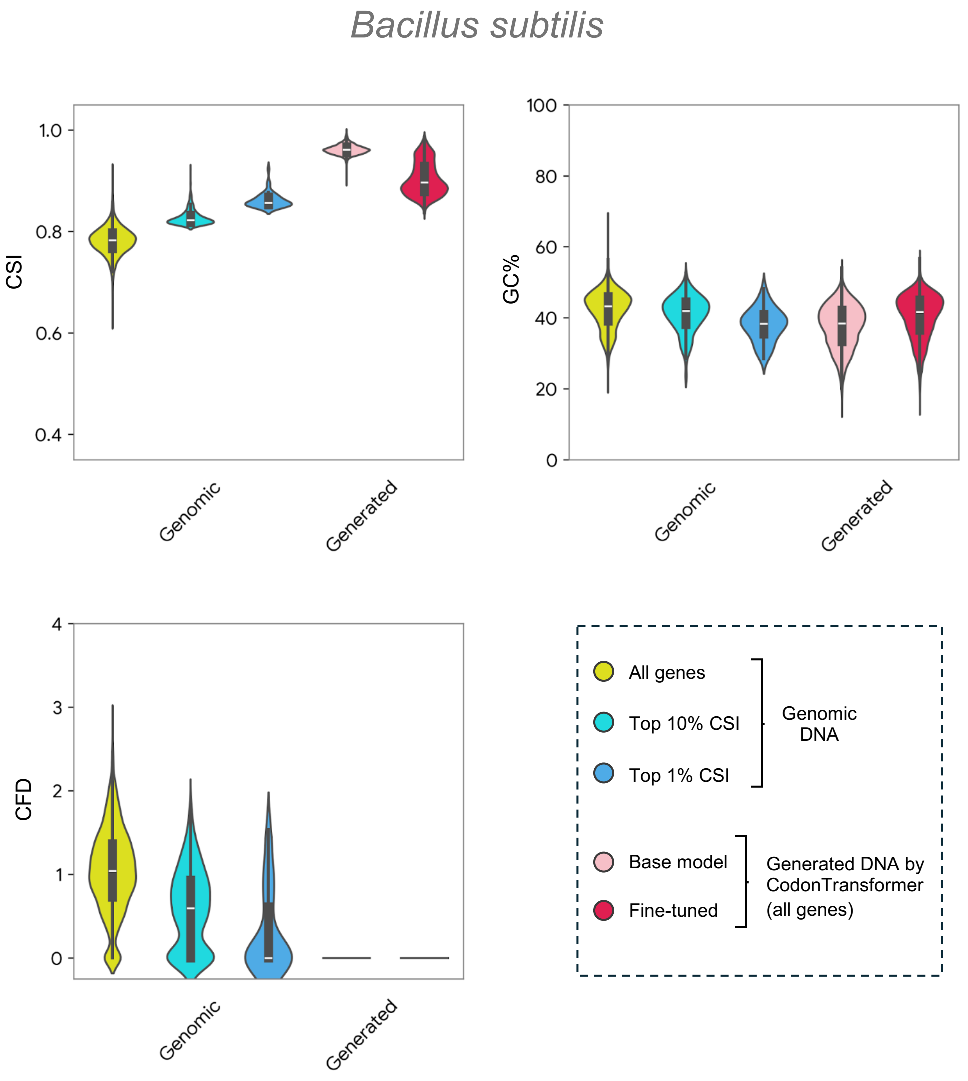


**Supplementary Fig. 3:** Codon similarity index (CSI), GC content, and codon frequency distribution (CFD) for genomic DNA sequences of *B. subtilis* and their generated counterparts by the base and fine-tuned CodonTransformer.

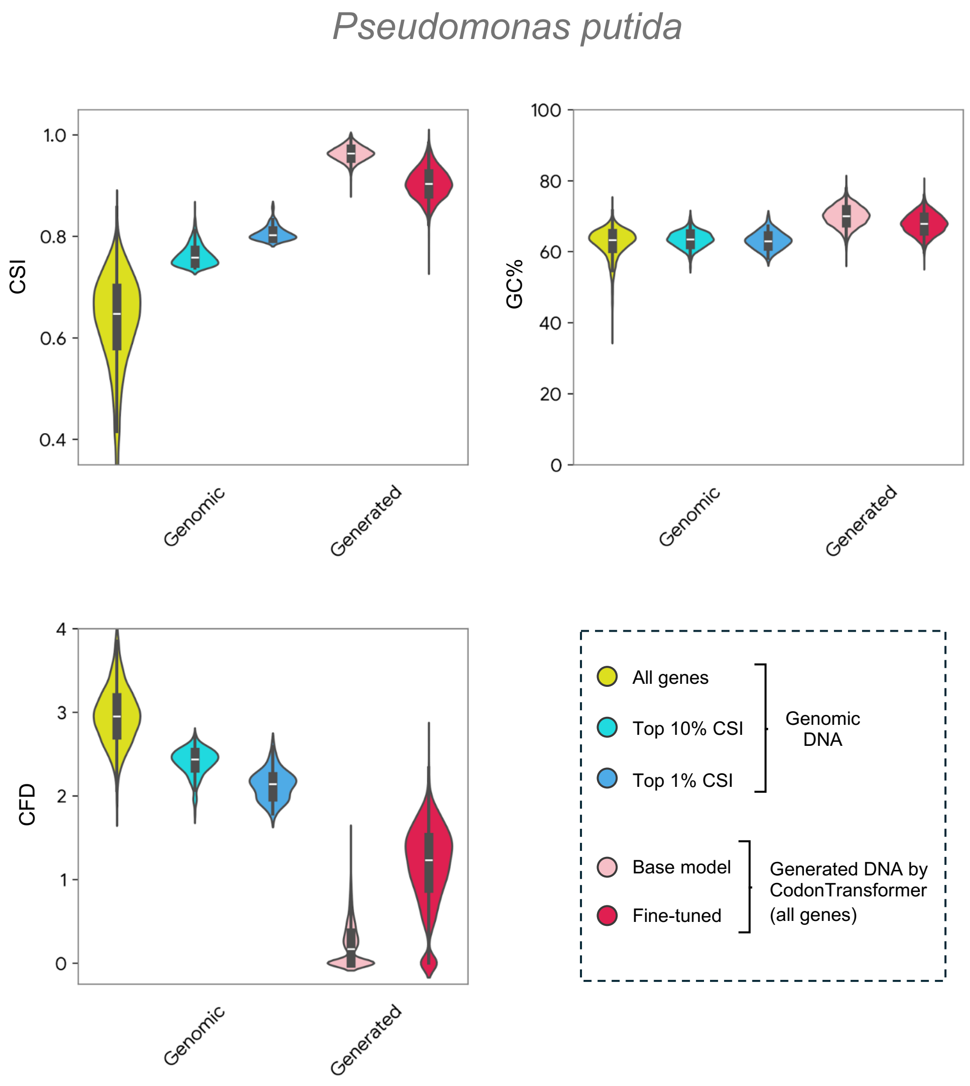


**Supplementary Fig. 4:** Codon similarity index (CSI), GC content, and codon frequency distribution (CFD) for genomic DNA sequences of *P. putida* and their generated counterparts by the base and fine-tuned CodonTransformer.

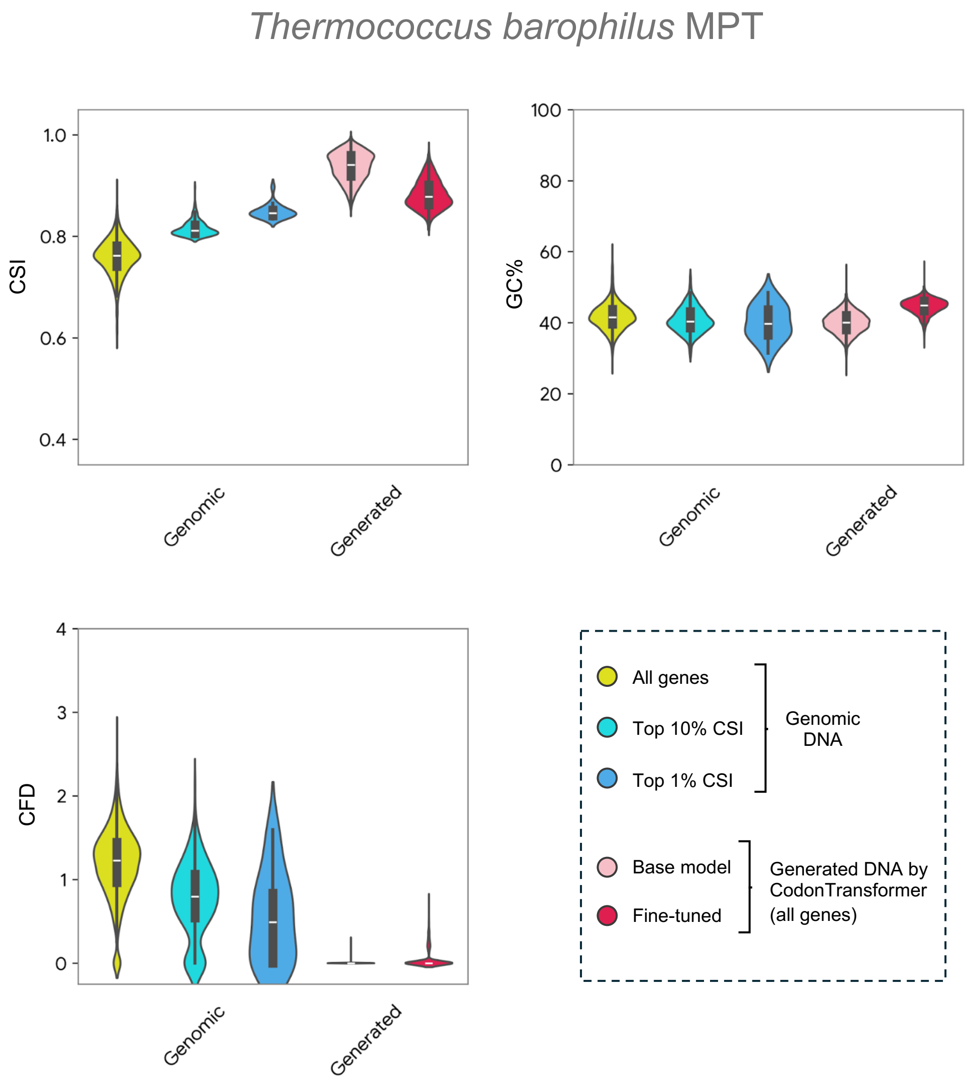


**Supplementary Fig. 5:** Codon similarity index (CSI), GC content, and codon frequency distribution (CFD) for genomic DNA sequences of *T. barophilus* and their generated counterparts by the base and fine-tuned CodonTransformer.

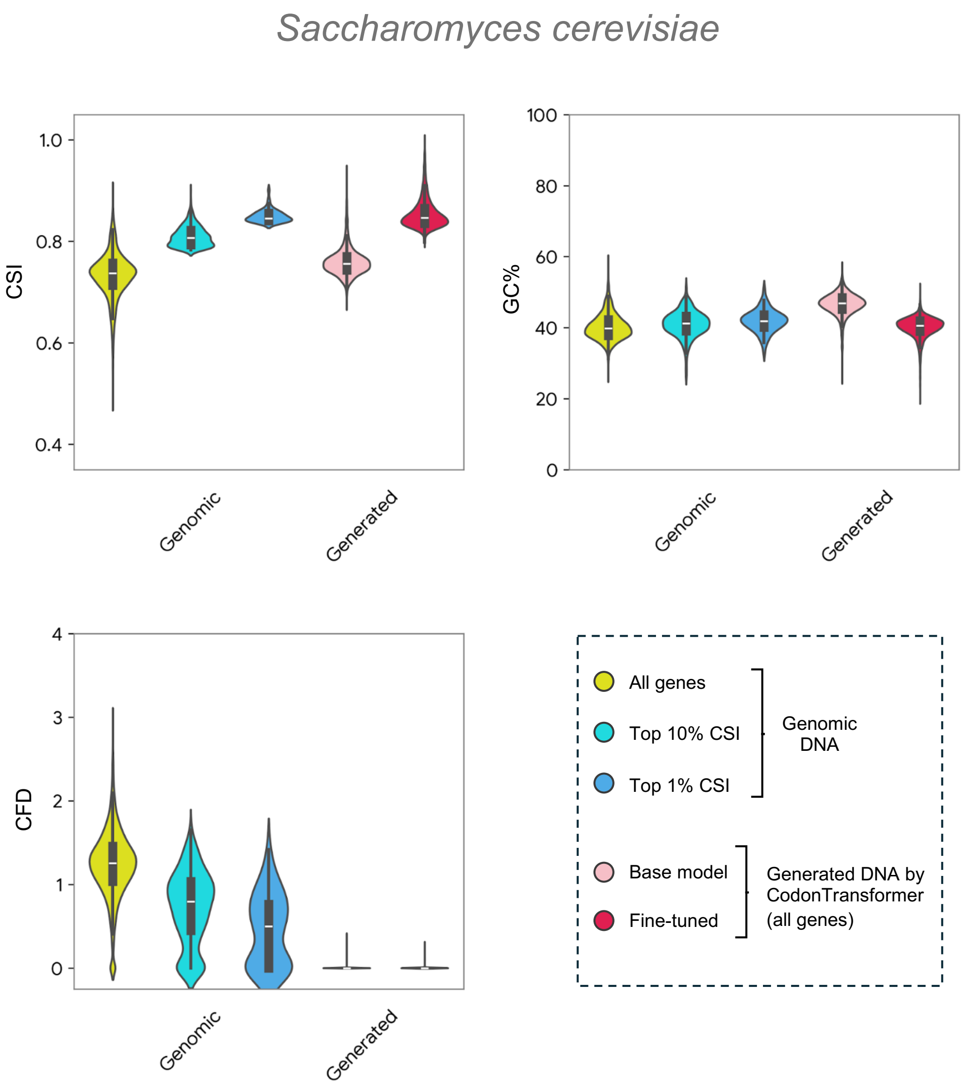


**Supplementary Fig. 6:** Codon similarity index (CSI), GC content, and codon frequency distribution (CFD) for genomic DNA sequences of *S. cerevisiae* and their generated counterparts by the base and fine-tuned CodonTransformer.

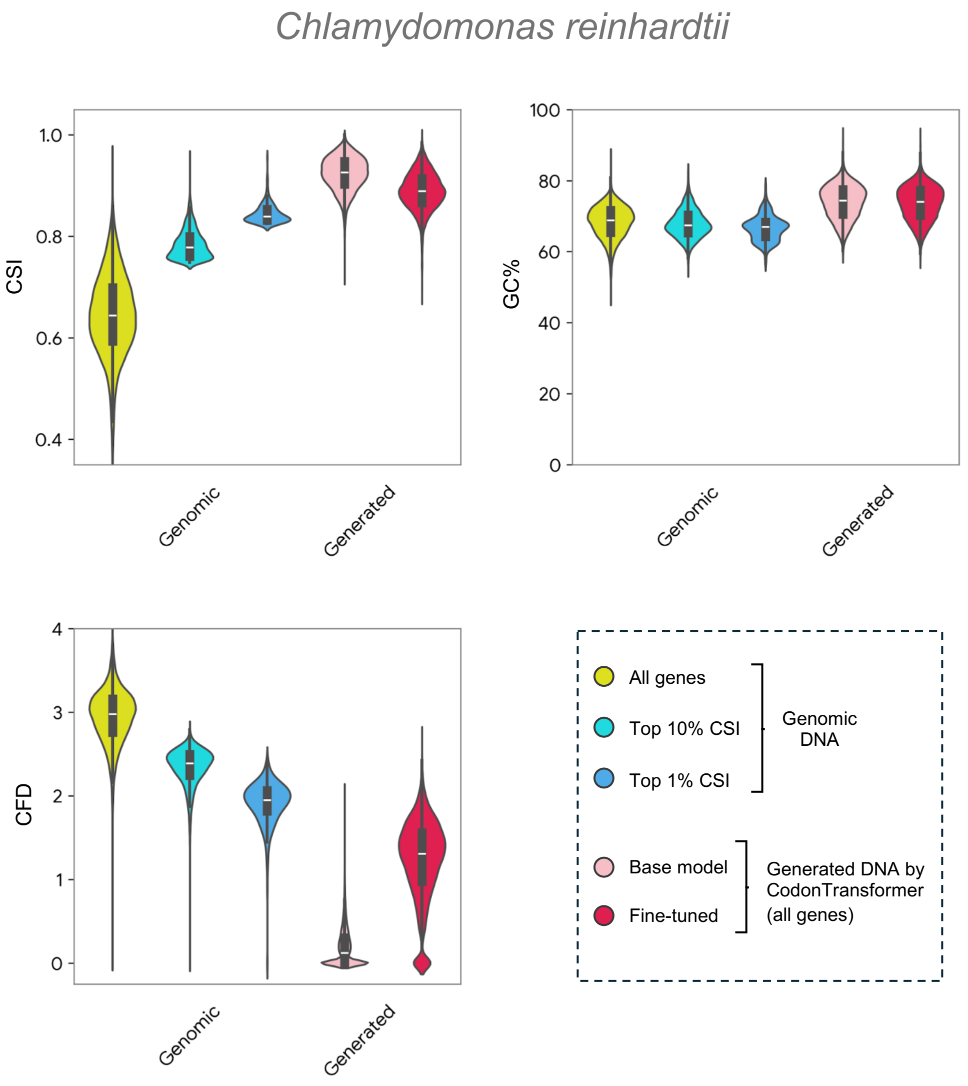


**Supplementary Fig. 7:** Codon similarity index (CSI), GC content, and codon frequency distribution (CFD) for genomic DNA sequences of *C. reinhardtii* and their generated counterparts by the base and fine-tuned CodonTransformer.

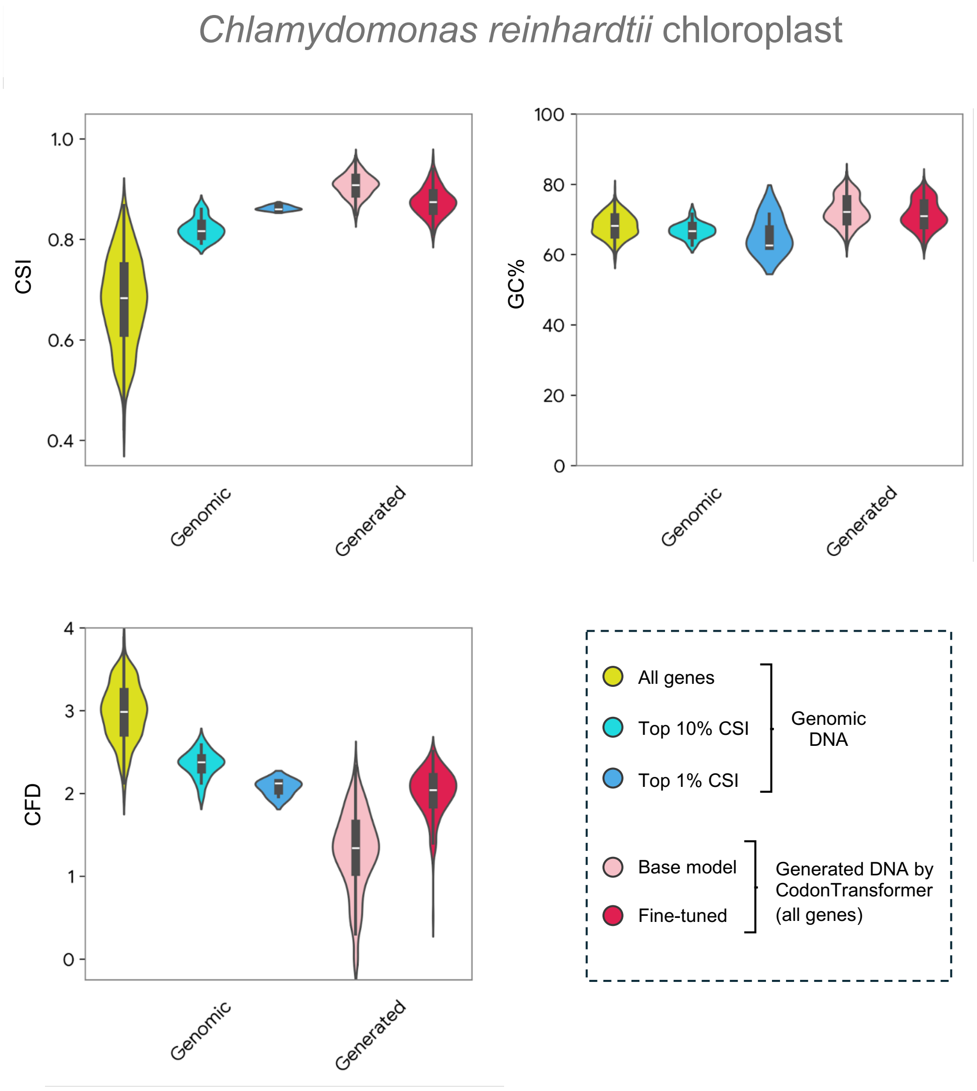


**Supplementary Fig. 8:** Codon similarity index (CSI), GC content, and codon frequency distribution (CFD) for genomic DNA sequences of *C. reinhardtii* chloroplast and their generated counterparts by the base and fine-tuned CodonTransformer.

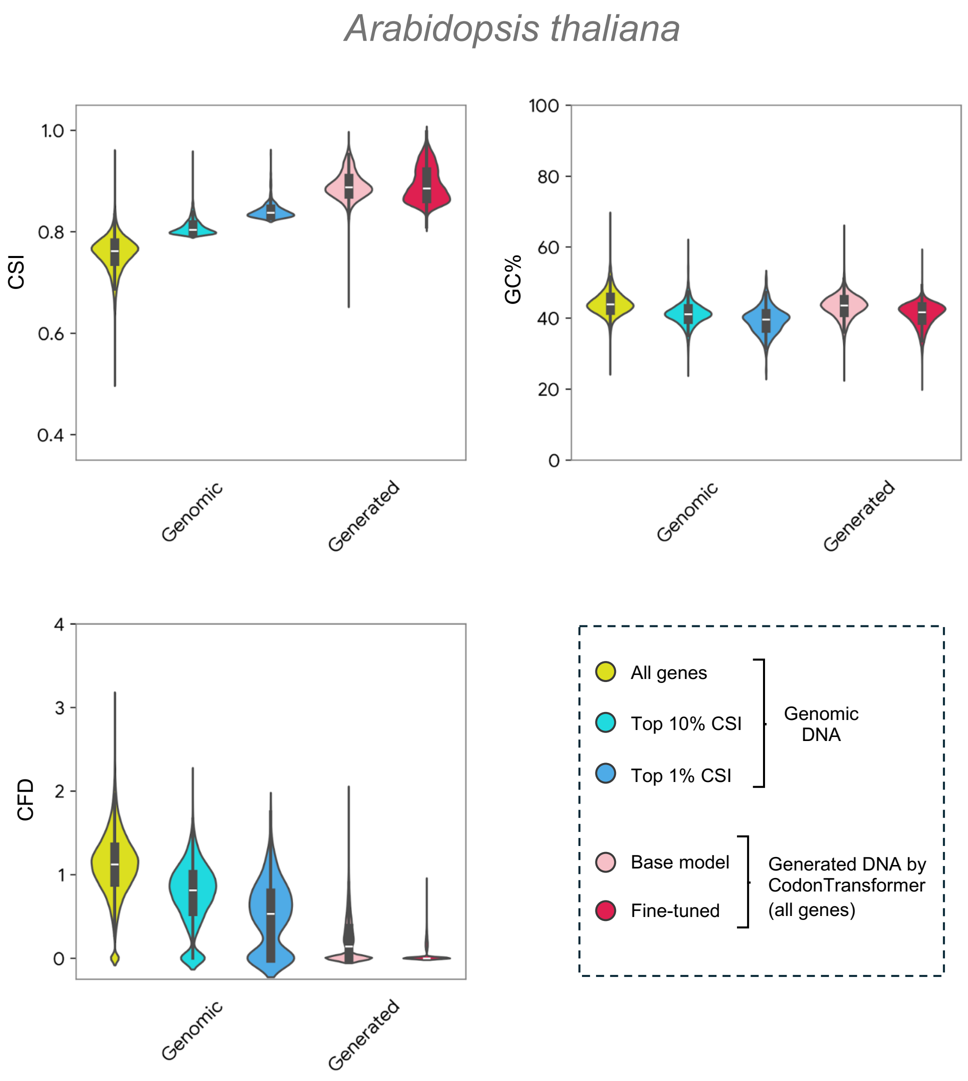


**Supplementary Fig. 9:** Codon similarity index (CSI), GC content, and codon frequency distribution (CFD) for genomic DNA sequences of *A. thaliana* and their generated counterparts by the base and fine-tuned CodonTransformer.

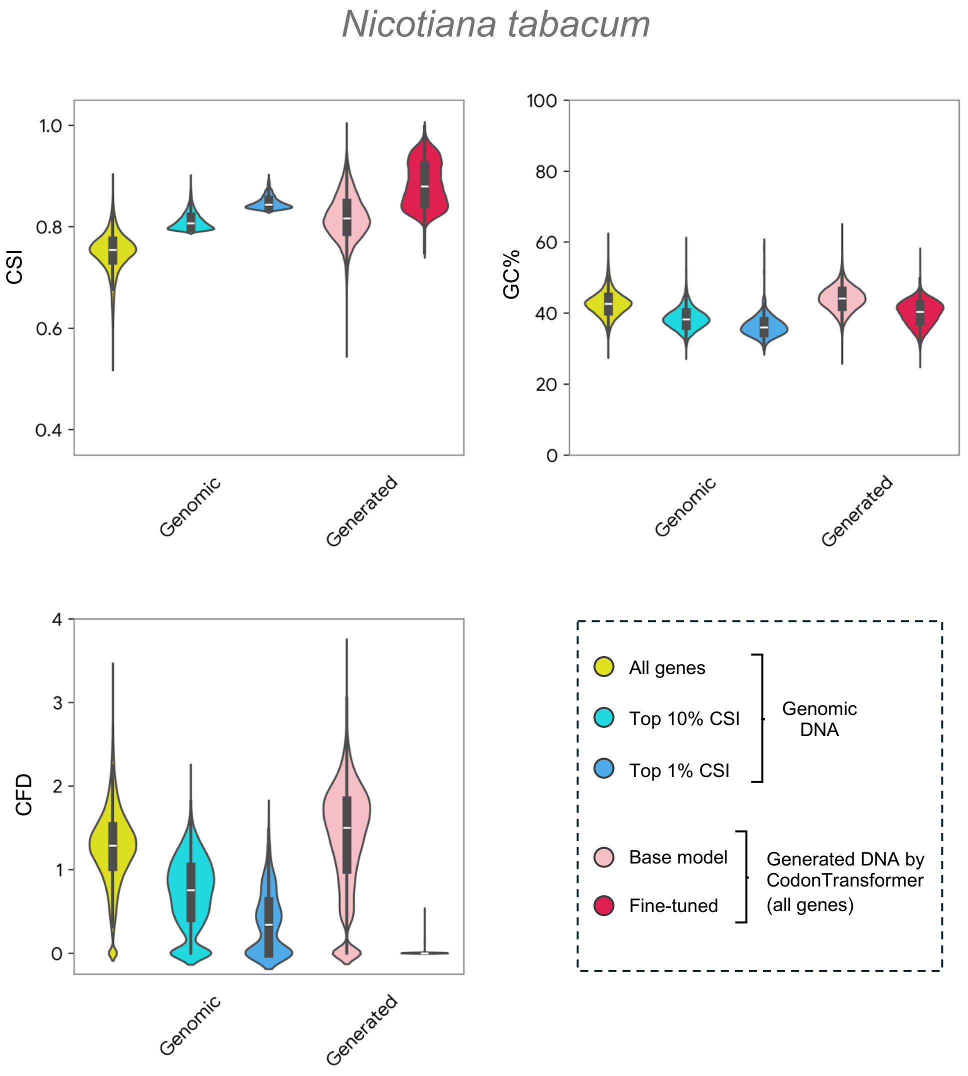


**Supplementary Fig. 10:** Codon similarity index (CSI), GC content, and codon frequency distribution (CFD) for genomic DNA sequences of *N. tabacum* and their generated counterparts by the base and fine-tuned CodonTransformer.

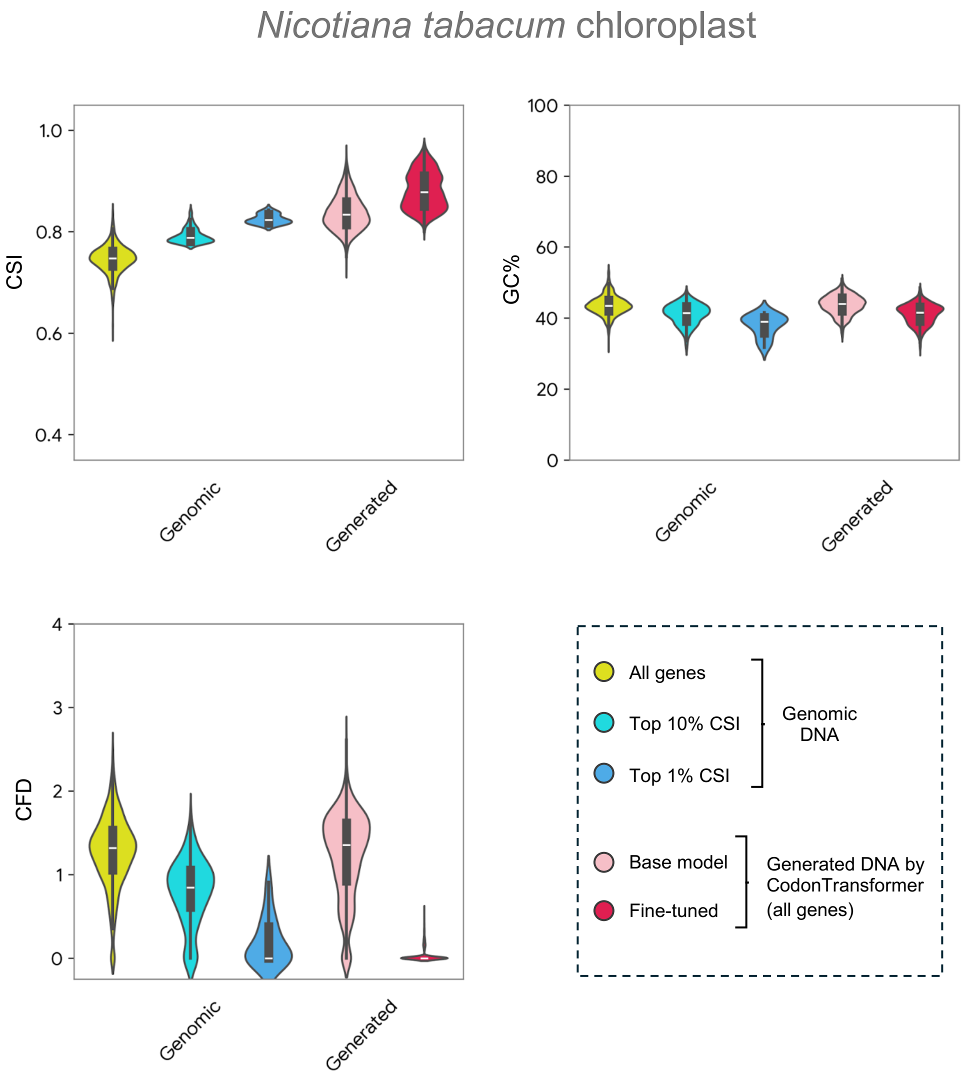


**Supplementary Fig. 11:** Codon similarity index (CSI), GC content, and codon frequency distribution (CFD) for genomic DNA sequences of *N. tabacum* chloroplast and their generated counterparts by the base and fine-tuned CodonTransformer.

*Caenorhabditis elegans*

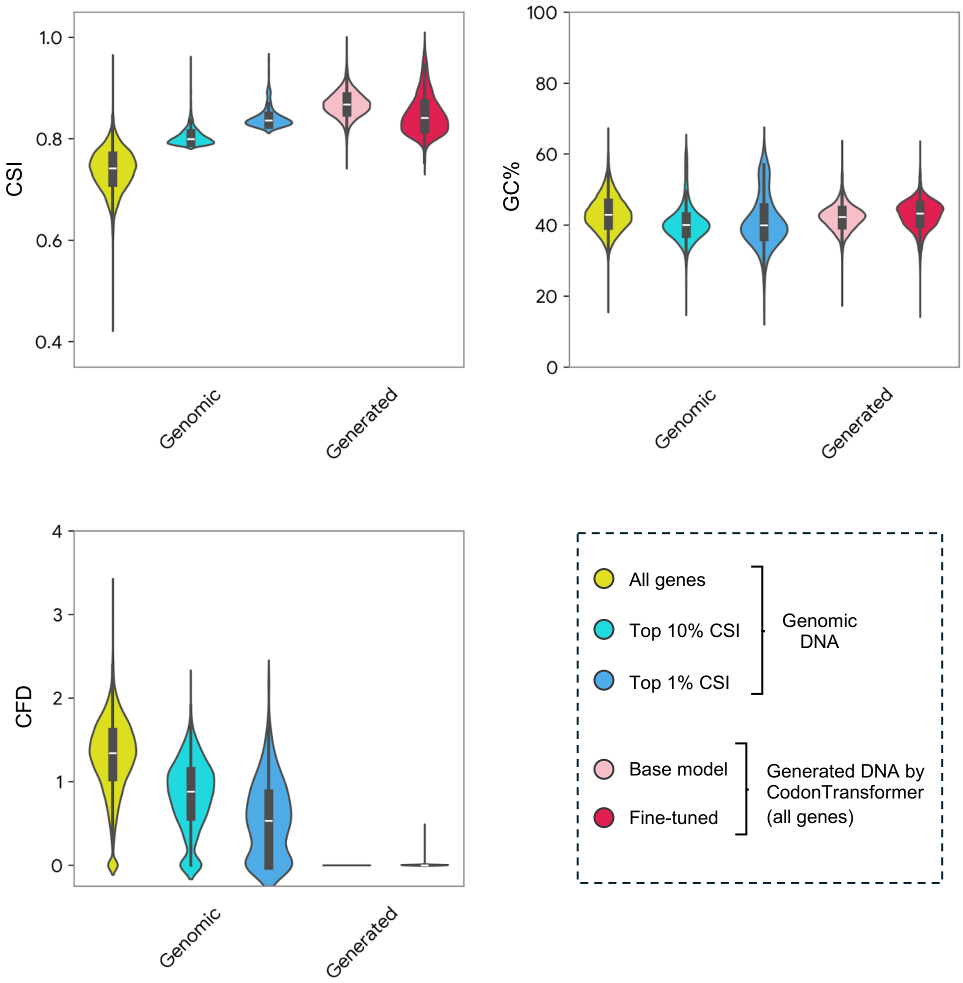


**Supplementary Fig. 12:** Codon similarity index (CSI), GC content, and codon frequency distribution (CFD) for genomic DNA sequences of *C. elegans* and their generated counterparts by the base and fine-tuned CodonTransformer.

*Danio rerio*

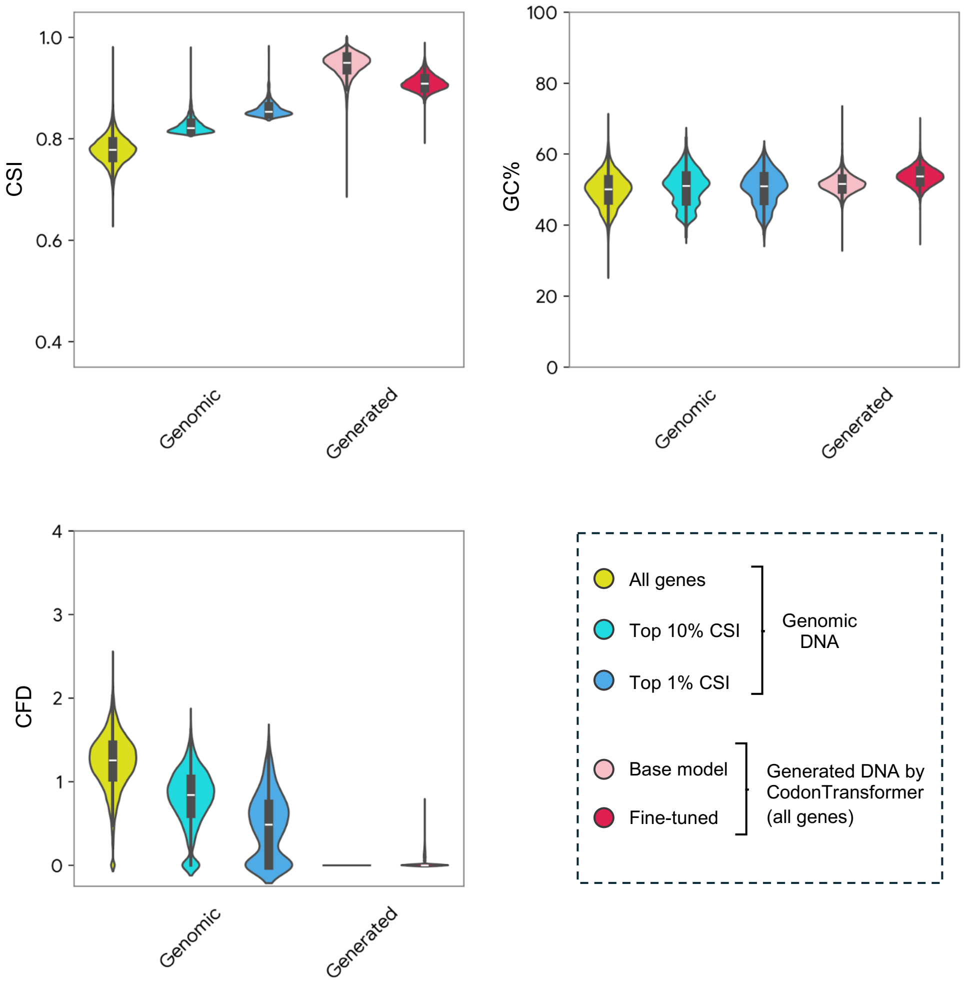


**Supplementary Fig. 13:** Codon similarity index (CSI), GC content, and codon frequency distribution (CFD) for genomic DNA sequences of *D. rerio* and their generated counterparts by the base and fine-tuned CodonTransformer.

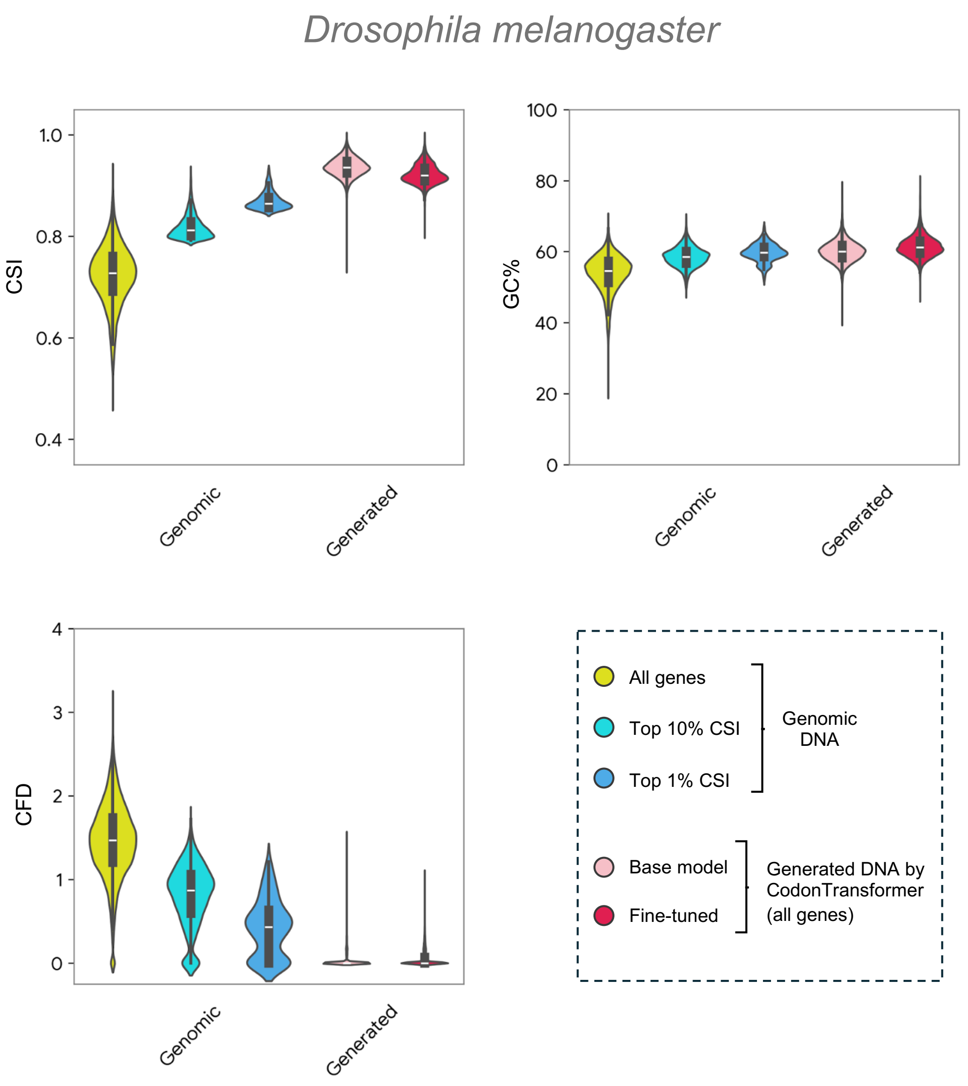


**Supplementary Fig. 14:** Codon similarity index (CSI), GC content, and codon frequency distribution (CFD) for genomic DNA sequences of *D. melanogaster* and their generated counterparts by the base and fine-tuned CodonTransformer.

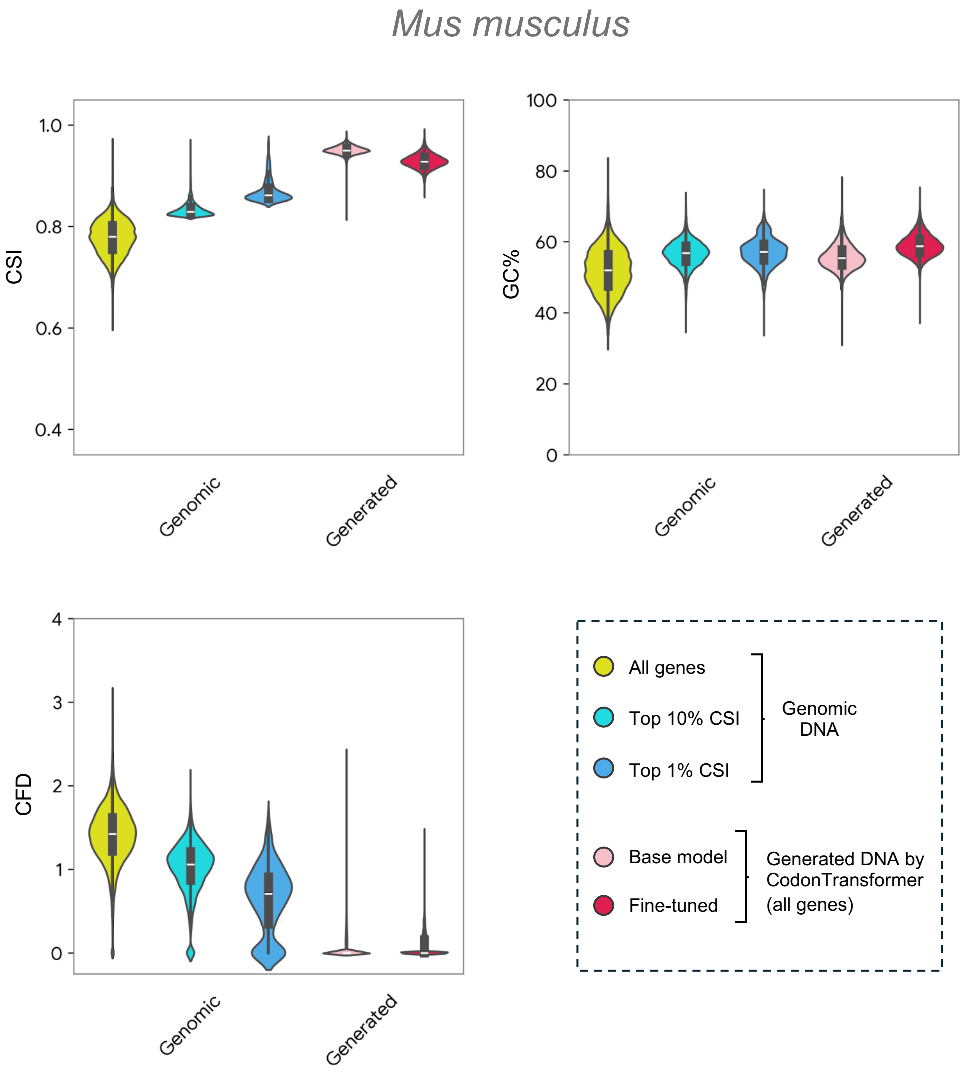


**Supplementary Fig. 15:** Codon similarity index (CSI), GC content, and codon frequency distribution (CFD) for genomic DNA sequences of *M. musculus* and their generated counterparts by the base and fine-tuned CodonTransformer.

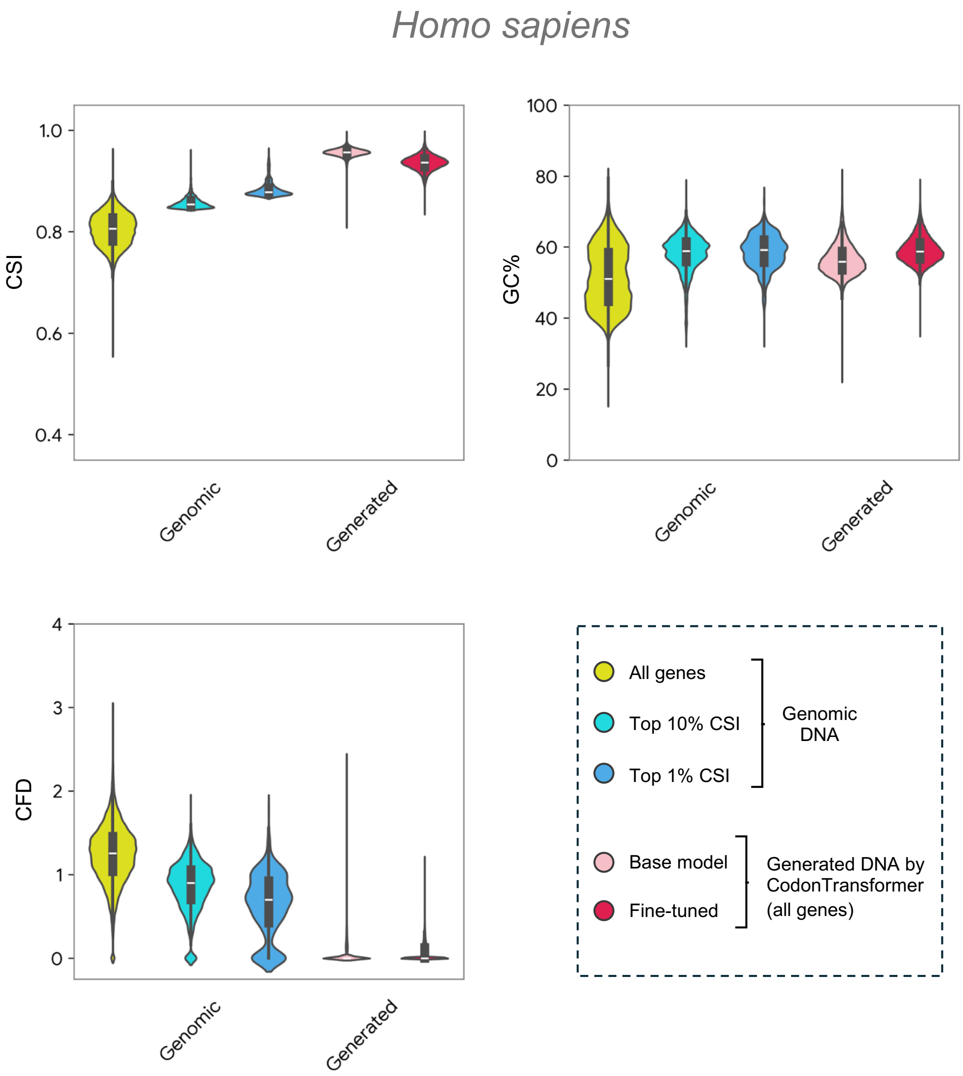


**Supplementary Fig. 16:** Codon similarity index (CSI), GC content, and codon frequency distribution (CFD) for genomic DNA sequences of *H. sapiens* and their generated counterparts by the base and fine-tuned CodonTransformer.

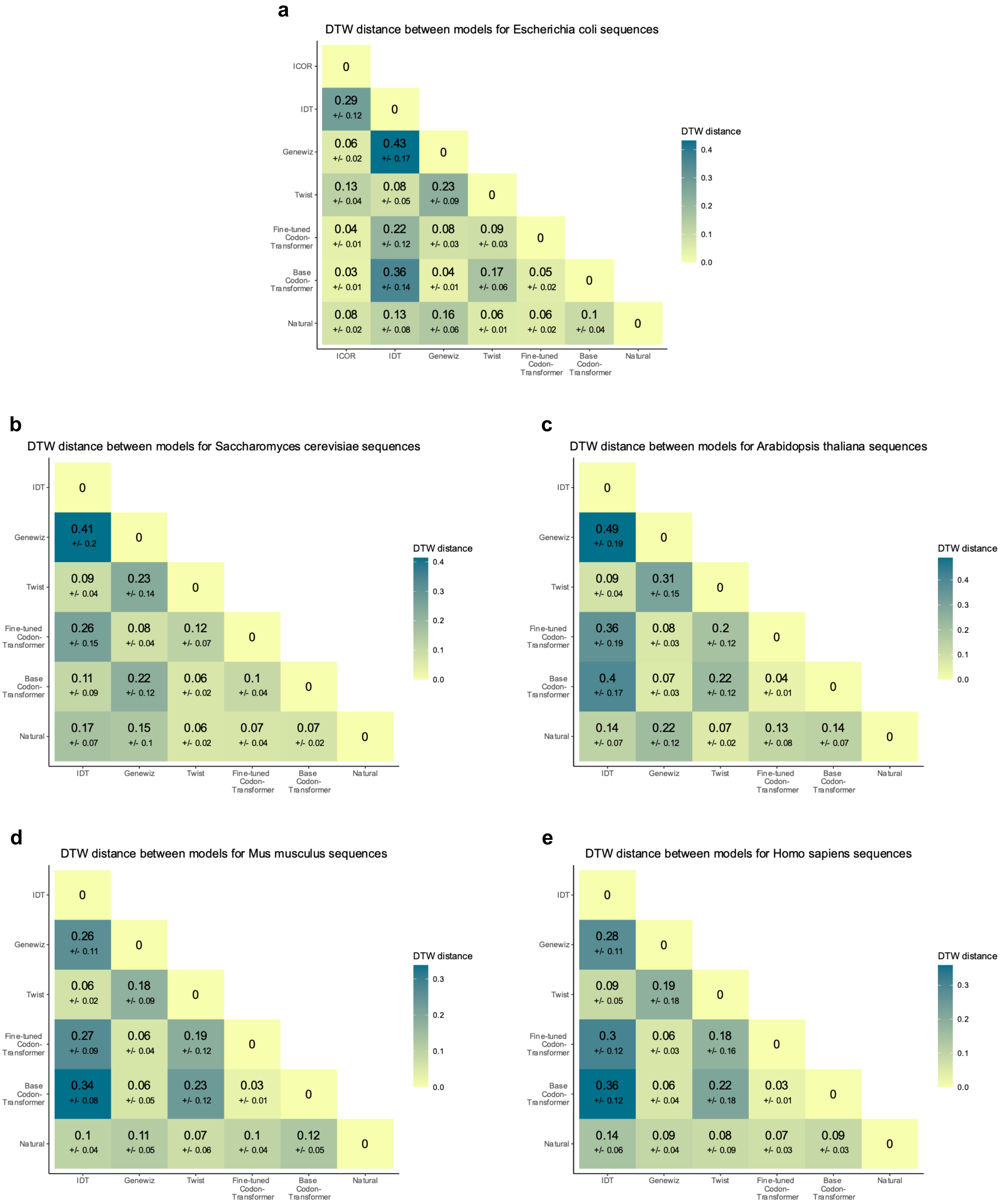


**Supplementary Fig. 17:** Model comparison based on normalized DTW distances between sequences generated for 50 random genes selected among top 10% CSI. Mean and standard deviation of normalized DTW distance between corresponding genes for *E. coli* (**a**), *S. cerevisiae* (**b**), *A. thaliana* (**c**), *M. musculus* (**d**), *H. sapiens* (**e**). Data underlying this figure is provided in **Supplementary Data 1**.

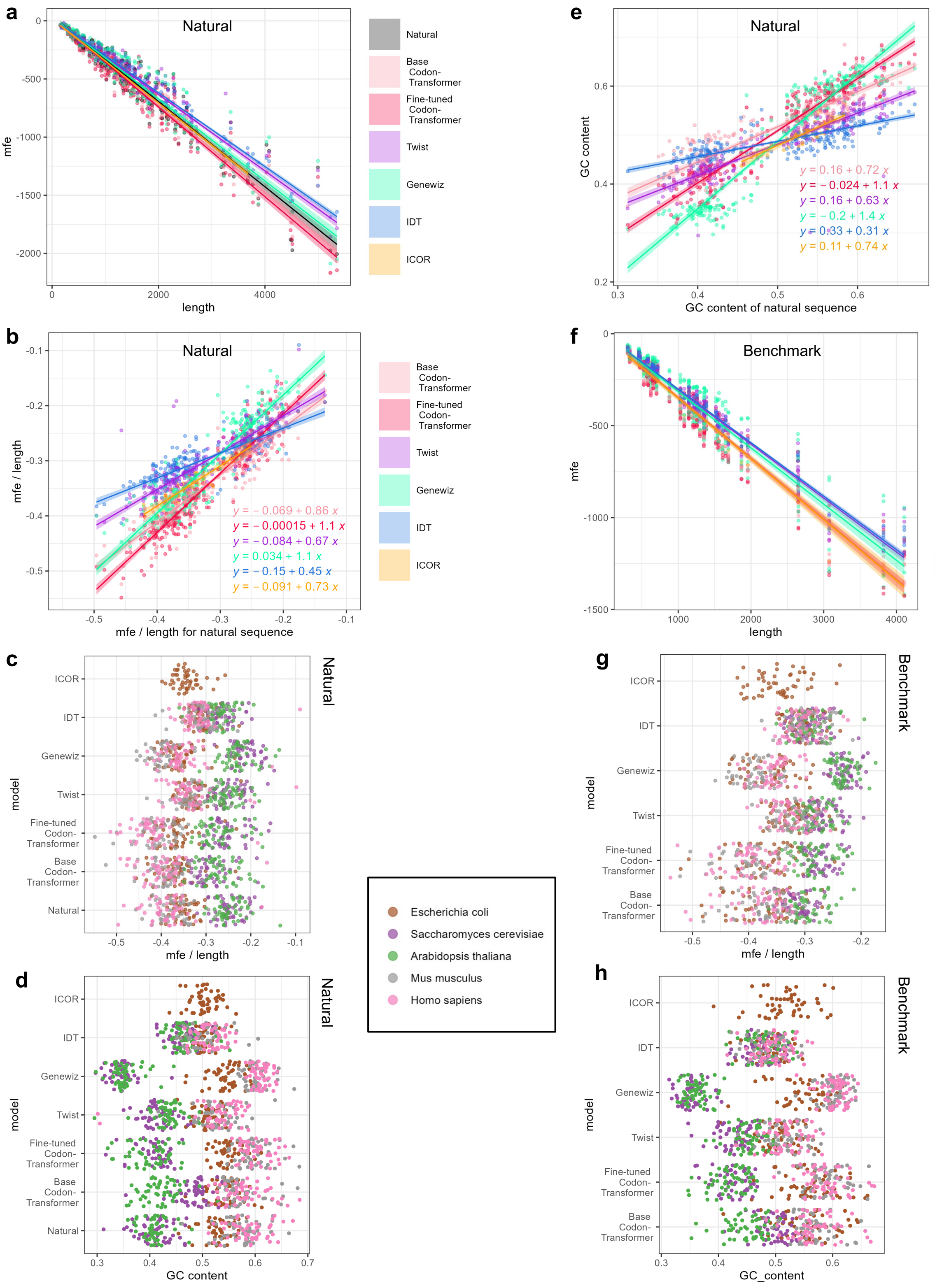


**Supplementary Fig. 18:** Model comparison based on minimum free energy (mfe) of RNA folding. Relationship between minimum folding energy and length for sequences generated by different models for 50 random genes among the top 10% CSI of each organism (**a**) and 52 benchmark proteins (**f**) for *E. coli, S. cerevisiae, A. thaliana, M. musculus, and H. sapiens*. **b**, Relationship between minimum folding energy normalized by protein length for generated and natural RNA. Minimum energy of RNA normalized by length for natural and benchmark proteins (**c** and **g**, respectively) and their GC content (**d** and **h**, respectively). **e**, Relationship between GC content of generated and natural. Data for **a-e** and **f-h** is provided in **Supplementary Data 1** and **2**, respectively.

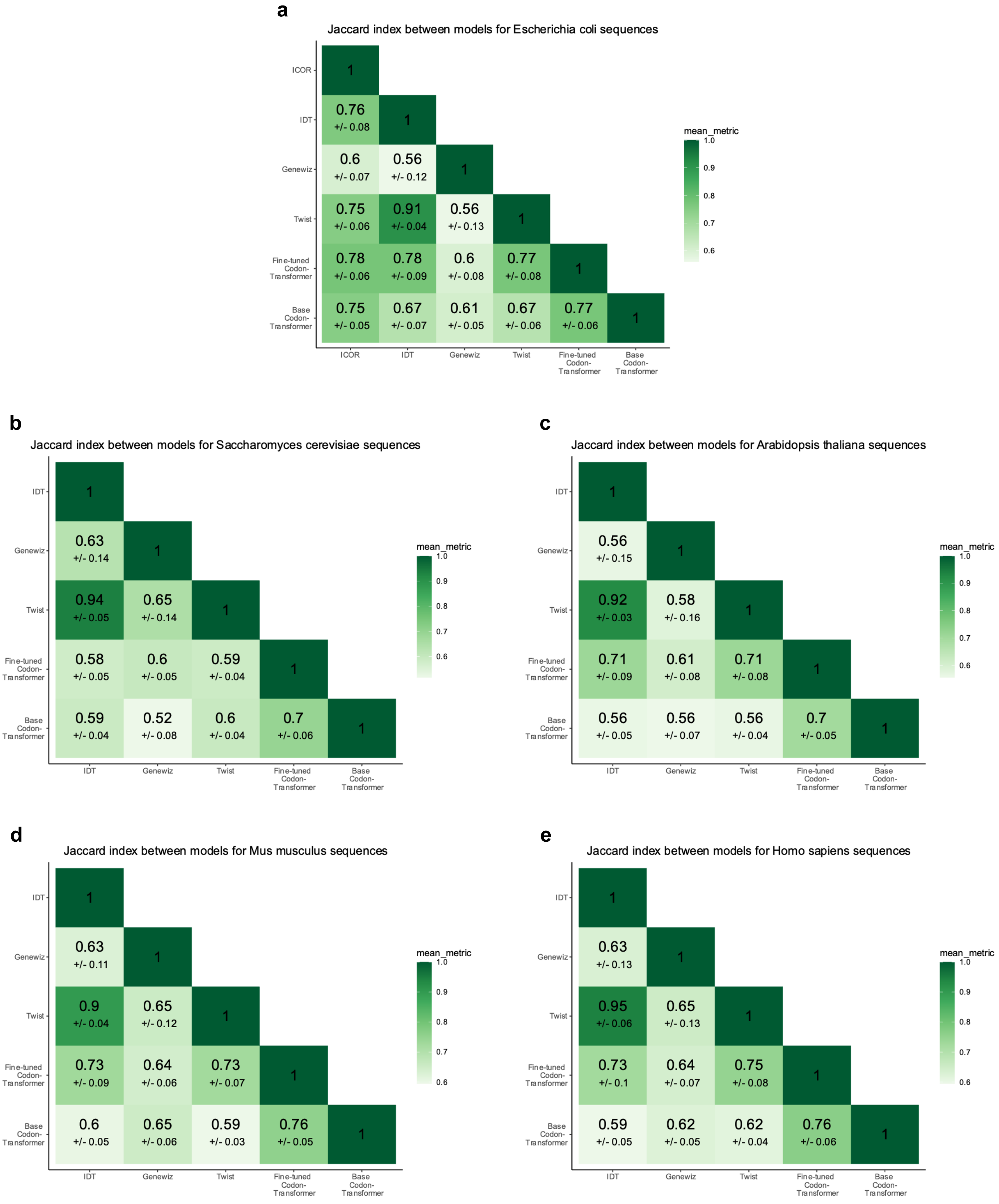


**Supplementary Fig. 19:** Model comparison based on Jaccard index between sequences generated for 52 benchmark proteins. Mean and standard deviation of Jaccard index between corresponding proteins for *E. coli* (**a**), *S. cerevisiae* (**b**), *A. thaliana* (**c**), *M. musculus* (**d**), *H. sapiens* (**e**). Data underlying this figure is provided in **Supplementary Data 2**.

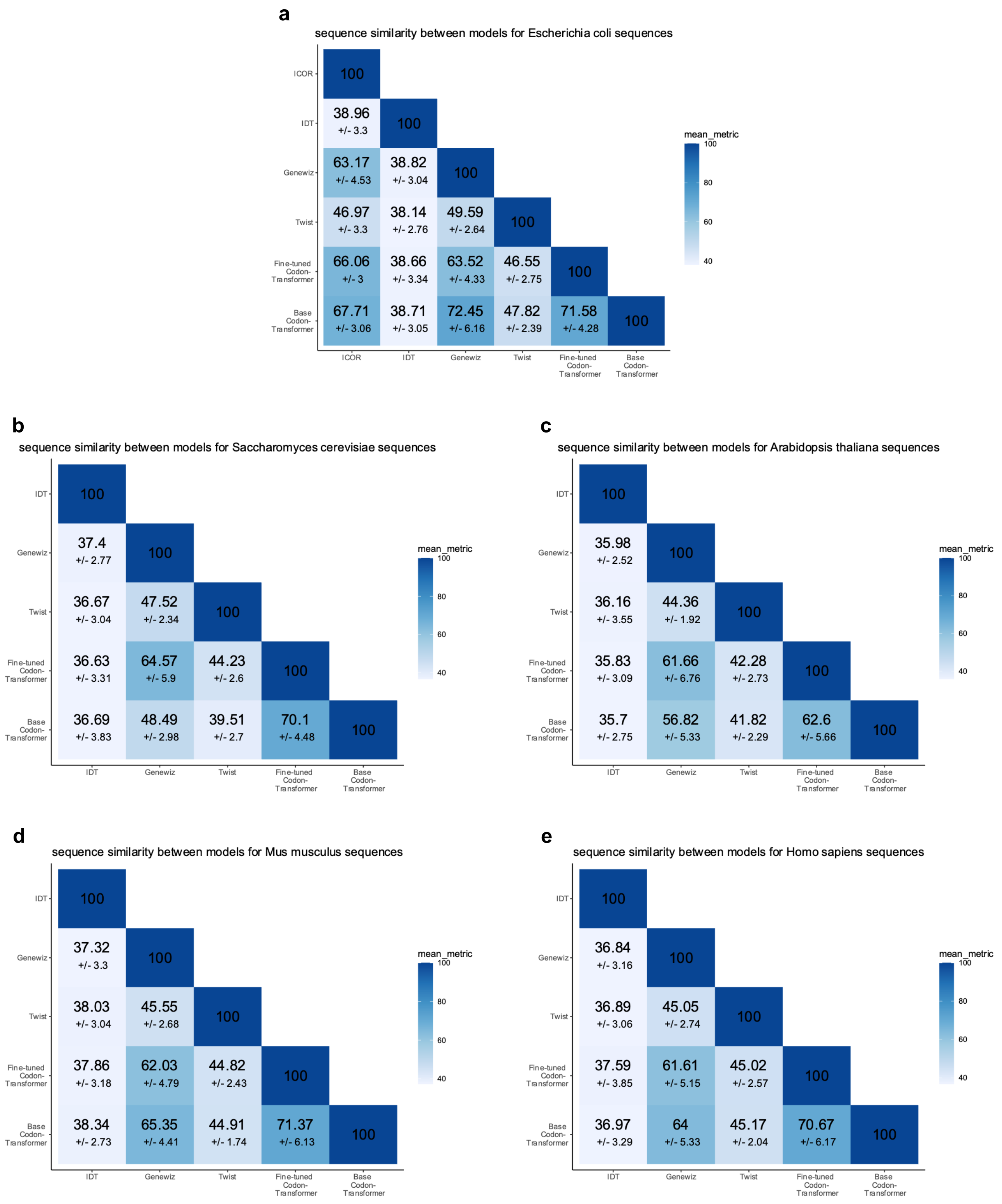


**Supplementary Fig. 20:** Model comparison based on sequence similarity between sequences generated for 52 benchmark proteins. Mean and standard deviation of sequence similarity between corresponding proteins for *E. coli* (**a**), *S. cerevisiae* (**b**), *A. thaliana* (**c**), *M. musculus* (**d**), *H. sapiens* (**e**). Data underlying this figure is provided in **Supplementary Data 2**.

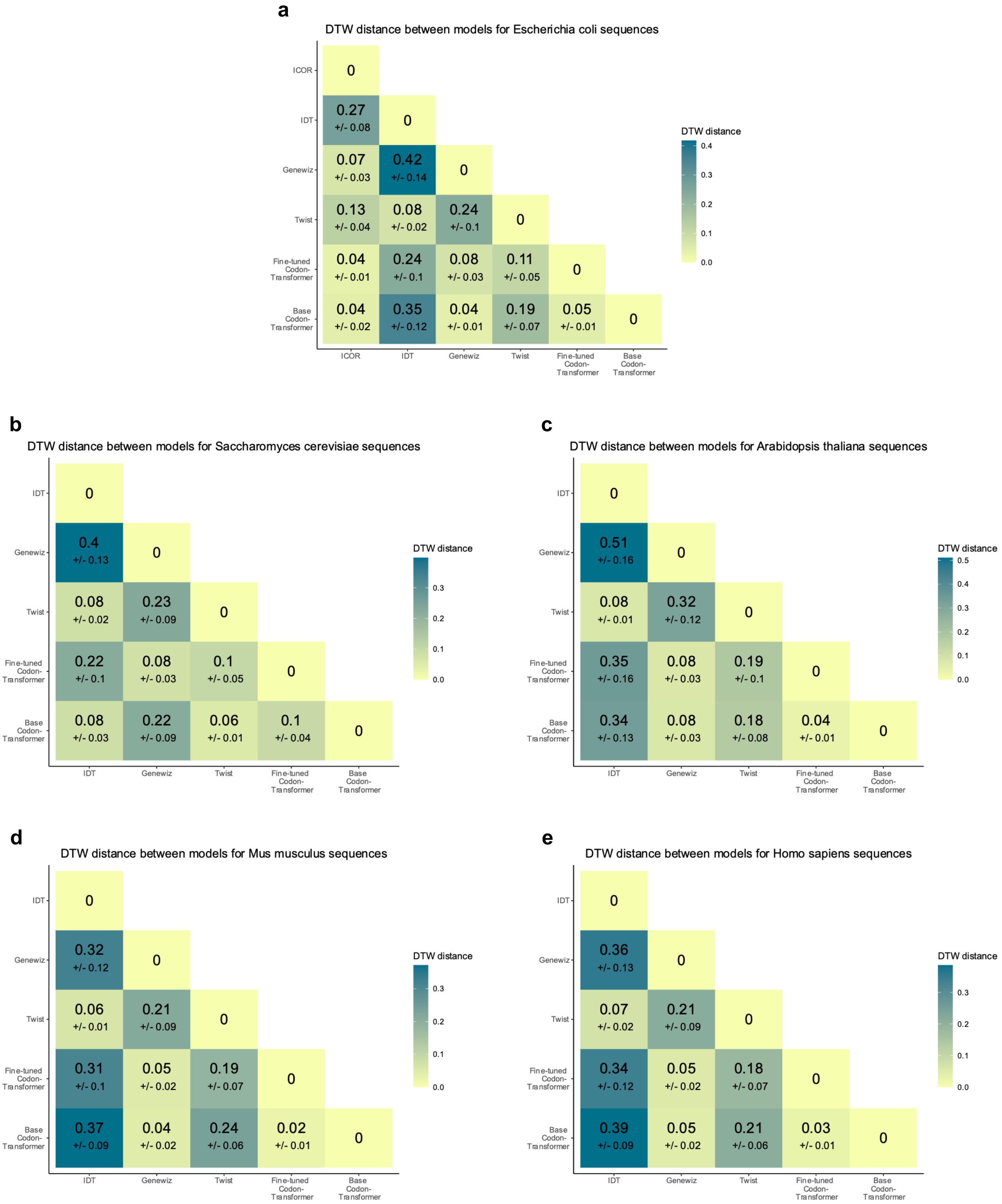


**Supplementary Fig. 21:** Model comparison based on normalized DTW distances between sequences generated for 52 benchmark proteins. Mean and standard deviation of normalized DTW distances between corresponding proteins for *E. coli* (**a**), *S. cerevisiae* (**b**), *A. thaliana* (**c**), *M. musculus* (**d**), *H. sapiens* (**e**). Data underlying this figure is provided in **Supplementary Data 2**.

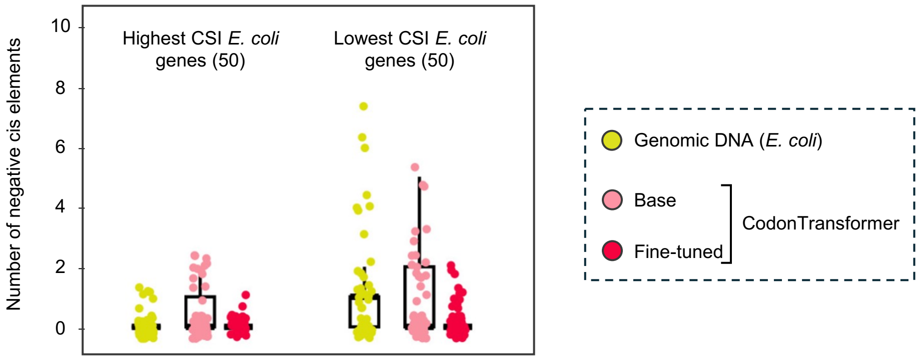


**Supplementary Fig. 22:** The average number of negative cis-regulatory elements of *E. coli* genes (from the general set), 50 lowest and 50 highest CSI, and for their codon-optimized sequences by the base and fine-tuned CodonTransformer**.**
